# Tag attachment reduces the initiation of recruitment in the rock ant, *Temnothorax rugatulus*

**DOI:** 10.1101/2023.08.19.553981

**Authors:** Benjamin Z. Taylor, Supraja Rajagopal, Takao Sasaki

## Abstract

Technological advances continue to push the boundaries of scientific inquiry in animal behavior. One such development is the emergence of automated tracking systems, which enable the collection of high-resolution spatio-temporal information for animals. Although tag-based tracking systems provide valuable insights into animal movement and collective behavior, the attachment of devices can have detrimental effects in some cases. Here, we investigated the effects of recently developed miniature tracking tags using the rock ant, *Temnothorax rugatulus*, as a model system. To do so, we compared the foraging activities of tagged and untagged ants within partially tagged colonies. Additionally, we compared the foraging activities of these tagged colonies with those of untagged control colonies. We found that tags did not reduce individual activity, with tagged ants visiting the food source as frequently as untagged ants. However, our analysis revealed a marked difference in recruitment behavior—tagged ants were significantly less likely to participate in tandem runs. This study demonstrates, for the first time, that tracking tags can negatively impact ant behavior. Although tracking devices are powerful tools for understanding complex behavioral patterns, it is crucial to carefully consider their potential impact on animal behavior to ensure accurate conclusions.

## Introduction

Technological advances continue to push the boundaries of scientific inquiry, allowing animal behavior researchers to obtain unprecedented datasets on the movement and interactions of animals [1, 2]. One such development is the emergence of automated tracking systems, which enable the collection of high-resolution spatio-temporal information [3, 4]. For instance, by attaching GPS tags to individual animals, researchers have gained insight into their movement patterns and social interactions during critical activities such as foraging [5, 6] and migrations [7]. Although tag-based tracking systems provide valuable insights into animal movement and collective behavior [1, 2, 8], research has shown that the attachment of devices can have detrimental effects in some cases [9], such as for birds [10] and bats [11]. Consequently, these adverse effects can introduce biases that compromise the accuracy of conclusions derived from the collected data [12].

Myrmecologists have recently started automating the acquisition of individual positional data for entire ant colonies (*ca*. 120 workers) over extended periods using miniature tags. These tags are attached to each ant and filmed using high-resolution cameras. The positions of these tags in each image are then automatically recorded using tracking open-source software built with pattern recognition libraries [13-15]. By analyzing these high spatio-temporal resolution data, researchers have revealed collective processes of ant colonies, such as emergent processes of social organization from workers [16], strategies for mitigating disease transmission within a colony [17], and mechanisms for spatial division of labor among workers [18]. Although automated tag-based tracking is becoming more commonly used to study ants, few have directly explored the impact of tag attachment on behavior.

To investigate tag effects on ant behavior, we used the rock ant, *Temnothorax rugatulus*, whose relatively small colony size (50-200 workers) and robustness to laboratory conditions have made them a model system for collective decision-making research [19-22]. Collective decisions rely on active scouts that find and assess options (*e*.*g*., candidate nest sites and food sources) and recruit nestmates with a probability dependent on option quality [22, 23]. When foraging, *Temnothorax* ants recruit via tandem runs, in which a knowledgeable leader directly guides a nestmate to the food source [24, 25]. In this study, we compare the foraging activity of tagged and untagged ants and test if tags impact the likelihood of 1) visiting food sources, 2) leading tandem runs, and 3) following tandem leaders.

## Methods

### Subjects

Six colonies of *T. rugatulus* were collected from rock crevices on Mount Lemmon (32°23’44.4”N 110°41’27.1”W) north of Tucson, Arizona, in July 2021 and October 2022. Each colony used in this study had at least one queen, several brood items, and 81 ± 12 workers (see Table 1 for colony details). In the lab, we housed colonies in artificial nests comprised of a cardboard sheet (2 mm thick) wedged between a pair of glass microscope slides (50×75 mm) [21]. Each cardboard sheet was cut to have a rectangular cavity (28×40 mm) and a single connecting entrance tunnel (15×4 mm) [22].

**Table 1.** Colony composition and the effect of tags on foraging behaviors.

|  | Colony | Whole colony |  | Feeder visit |  |  | Tandem-running leader |  |  | Tandem-running follower |  |  |
| --- | --- | --- | --- | --- | --- | --- | --- | --- | --- | --- | --- | --- |
|  |  | Tagged | Untagged | Tagged | Untagged | P value | Tagged | Untagged | P value | Tagged | Untagged | P value |
| Tagged colonies | R2119 | 61 (95%) | 3 (5%) | 35 (92%) | 3 (8%) | 0.26 | 0 | 0 | – | 0 | 0 | – |
|  | R2111 | 62 (69%) | 28 (31%) | 36 (56%) | 28 (44%) | **< 0.001** | 0 (0%) | 3 (100%) | **0.03** | 2 (67%) | 1 (33%) | 1.0 |
|  | R2100 | 93 (95%) | 5 (5%) | 22 (100%) | 0 (0%) | 0.58 | 0 | 0 | – | 0 | 0 | – |
| Untagged colonies | FR2217 | – | 68 (100%) | – | 55 (100%) | – | – | 10 (100%) | – | – | 10 (100%) | – |
|  | FR2218 | – | 83 (100%) | – | 51 (100%) | – | – | 3 (100%) | – | – | 3 (100%) | – |
|  | FR2220 | – | 71 (100%) | – | 143 (100%) | – | – | 12 (100%) | – | – | 12 (100%) | – |

The "Whole colony" column provides counts and proportions (%) of tagged and untagged workers for each colony, with a solid line indicating the separation between tagged colonies and untagged control colonies. The columns for "Feeder visit," "Tandem-running leader," and "Tandem-running follower" provide the total number of events observed for each behavior and, when applicable, p-values obtained from Fisher's exact test. Note that feeder visit counts exclude tandem-running leaders and followers.

Nests were kept in plastic boxes (11×11 cm) with the walls coated in Fluon to prevent the ants from escaping. Colonies were maintained in an incubator at 13°C on a 12:12 light/dark cycle and were provided with water, spam, and agar-based food [26] *ad libitum*.

### Tags

We generated unique two-dimensional matrix barcodes with the Python package Pinpoint [27]. Tags (0.8×0.8 mm) were printed on thin, durable paper (OK Kinfuji, SunM Color Co., Ltd., Japan) and weighed approximately 0.2 mg each. To attach the tag, we gently pushed an ant onto a soft, sticky surface (kneaded rubber; RDD-150, Sakura) and then secured her head with a staple. We then placed a drop (∼ 1 nl) of super glue (LPJ-005, Loctite®) on her thorax and placed the tag on it. Finally, the tagged ant was gently removed from the kneaded rubber after 6 minutes (allowing the glue to cure and the tag to firmly adhere to the ant) before returning her to the colony. We visually confirmed that all the tagged ants walked normally in the nest (Fig 1a & Fig 1b).

**Fig 1.**
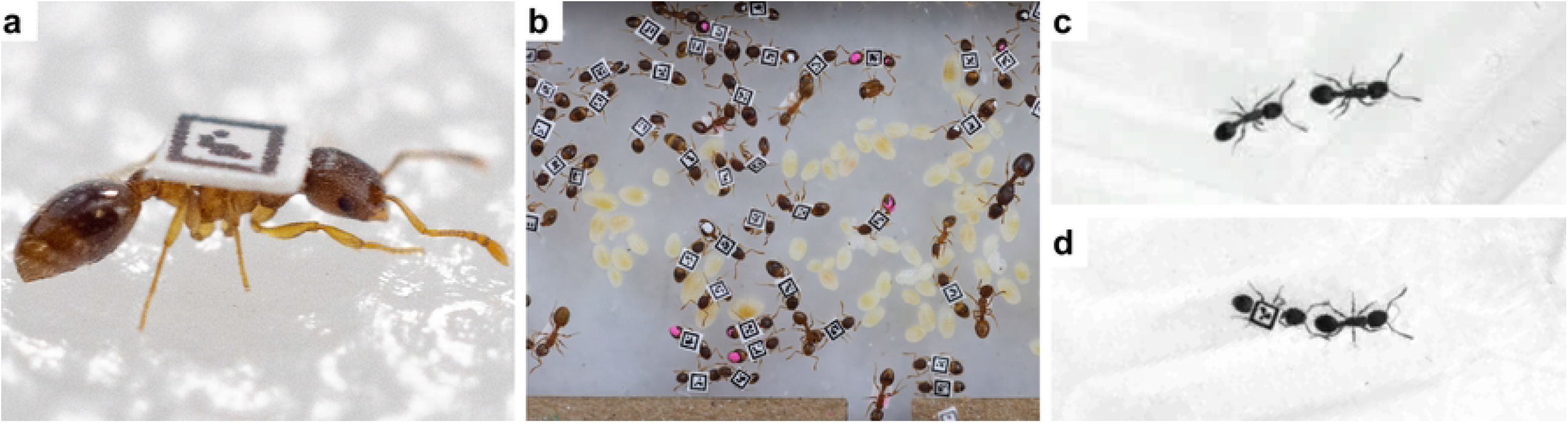
Tagged ants: (**a**) An ant (*T. rugatulus*) fit with a miniature QR tag (0.8×0.8 mm). (**b**) A colony of tagged workers inside a home nest. (**c**) Untagged scouts tandem running. (**d**) An untagged scout leads a tagged follower in a tandem run during a foraging experiment.

Workers in three colonies were tagged prior to foraging experiments. However, the final proportion of tagged individuals varied since tags were frequently detached during grooming and handling (Table 1). Three colonies were never tagged and served as control groups (“untagged colonies” in Table 1).

### Experimental Procedure

We conducted all experiments in a box arena (24×24 cm). Since *Temnothorax* ants tend to walk along edges [28], we lined the arena floor with a hexagonal array to promote route variations [29] (Fig 2c). At the beginning of the experiment, we placed a home nest containing a colony on one side of the arena and a feeder on the opposite side (Fig 2c). The feeder consisted of a glass depression slide (25×75 mm) with 0.2 ml of 0.8 mol sugar solution. We chose this concentration since *Temnothorax* ants show a strong recruitment effort towards it [22, 29]. Six high-resolution (7,716×5,360 pixels) cameras (DFK AFU420-CCS, The Imaging Source®) sufficient for reading miniaturized tags were suspended 16 cm above the arena (Fig 2b). Each camera captured a portion of the arena, jointly covering the entire arena. Immediately after placing the home nest and feeder in the arena, we started recording. We synchronized our six-camera system using Norpix StreamPix and recorded for approximately 6 hours. All colonies were starved for two weeks before the experiment to motivate foraging and only tested once. Before each experiment, the arena was cleaned with ethanol to remove all chemical marks.

**Fig 2.**
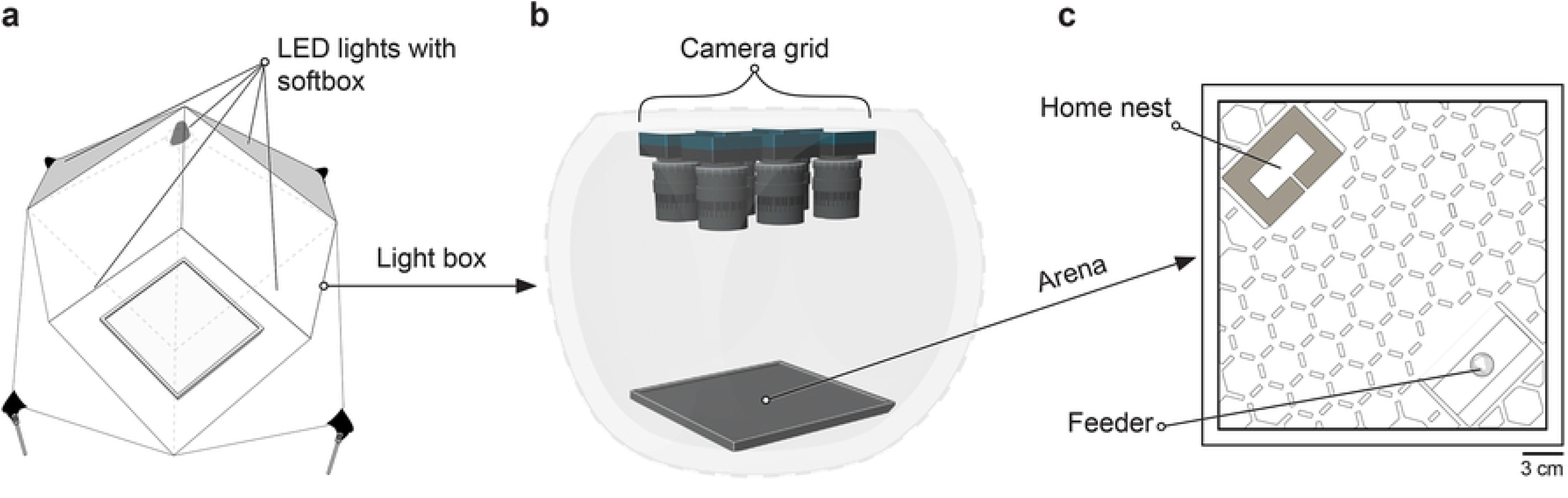
Multi-camera tracking system: (**a**) The system’s external perspective (aerial) with five LED light boxes surrounding a lightbox-enclosed arena (gray, dashed line). (**b**) The encased lightbox, containing the arena and the camera grid. (**c**) An aerial view of the box arena with the home nest opposite the feeder (food source).

### Video processing

In this experiment, the original high-resolution footage could be reduced since tags no longer needed to be machine-readable, just visible for their presence. Therefore, raw video dimensions were first scaled down 7.5 times using an open-source video editing software FFmpeg [30]. Since each camera captured only part of the arena, we then acquired transformation matrices to convert images to the same coordinate system and extend the field of view to the entire arena. These transformation matrices were determined and visually verified for our camera grid using a software package for ImageJ, BigStitcher [31]. We then batch-applied these transforms to our reduced image set with the Python library OpenCV [32], resulting in a low-resolution stitched video.

This procedure produced a low-resolution, stitched video file that can be easily watched and quickly transferred between computers without demanding a high-end GPU or ample disk space.

### Analysis

We watched 5 hours and 30 minutes of footage from each experiment (S1 Video Repository). We chose a significantly longer duration than previous foraging experiments of *Temnothorax* ants [22, 29] to ensure we captured all foraging activity. We manually annotated videos, recording the status of each ant (tagged or untagged) and time of occurrence for three foraging events: 1.) feeding at food source, 2.) leading a tandem run, and 3.) following a tandem run (Fig 1c, d). All statistical analyses were performed in Python (v. 3.9.13) with the SciPy [33] (v. 1.10.0) and Statsmodels [34] (v. 0.14.0) libraries.

We investigated the effects of tags on these events by examining the participation of tagged and untagged ants within tagged colonies. Most ants had tags in these colonies, with untagged ants comprising 5% for two colonies and 31% for the third colony (Table 1). We employed Fisher’s Exact tests to assess whether tagged and untagged ants exhibited equal participation in foraging activities relative to their presence in the colony. We chose this statistical method due to its suitability for analyzing small sample sizes and contingency tables with zero counts, which are common in our dataset.

To assess the overall impact of tags on colony-level foraging activity, we examined whether tagged colonies exhibited more feeder visits and/or tandem runs than untagged colonies using Welch’s t-test. Furthermore, to account for variations in colony composition of tagged and untagged ants, especially the number of feeder visits by these ants, we also employed ordinary least squares regression analysis using the number of tandem running as a dependent variable and (1) the number of untagged ants within the colony and (2) the number of feeder visits by untagged ants as two independent variables. These colony-level comparisons provide insights into the relationship between tags and tandem-running behavior, offering both direct evidence through comparisons of tagged and untagged colonies and indirect evidence by considering the underlying factors influencing tandem-running behavior.

## Results

### Tags had no effect on feeder visits

When we looked at the tagged colonies only, we found no significant relationship between tagged ants and their presence at the food source relative to the proportion of tagged ants in the colony for two of the three colonies (Table 1). When we compared tagged and untagged colonies, the frequency of feeder visits (relative to the colony size) was not significantly different between these colonies (Welch’s t-test, one-sided: T = 1.29, df = 4, *P* = 0.15). These data suggest that tags do not have a significant effect on the average number of feeder visits.

### Tags impact recruitment of nestmates

Among the three tagged colonies, we only observed tandem runs in one colony (R2111), with three pairs of tandem runs. Notably, all the leaders involved in these runs were untagged, while two of the three followers had tags (Table 1). Fisher’s exact tests confirmed a significant difference in the proportions of tagged and untagged ants leading tandem runs (Fisher exact test, two-sided: *P* = 0.03), indicating a relationship between tags and leading tandem runs. However, no significant difference was found in the proportions of tagged and untagged ants following tandem runs (Fisher exact test, two-sided: *P* = 1.00).

In our comparative analysis between tagged and untagged colonies, we found that tagged colonies demonstrated a significantly lower frequency of tandem runs than untagged ones (Welch’s t-test, one-sided: T = 2.52, df = 4, *P* = 0.05). This finding indicates an association between tagging and a reduced likelihood of initiating tandem runs.

Additionally, we examined the relationship between the total number of feeder visits and the occurrence of tandem runs. There was a nearly significant correlation between the total number of feeder visits and the number of tandem runs (S1 Table 1). However, when considering the comprising groups and tandem running events, untagged feeder visits were significantly correlated (Pearson’s correlation, two-sided: Coefficient = 0.89, *P* = 0.02), whereas tagged feeder visits were not significantly correlated (Pearson’s correlation, two-sided: Coefficient = -0.72, *P* = 0.11).

The positive linear correlation between the number of untagged ants visiting the feeder and the occurrence of tandem runs suggests that feeder visits by untagged ants are a reliable predictor of tandem run occurrence (Fig 3). In other words, as the number of untagged ants visiting the feeder increases, the likelihood of tandem runs also increases. Moreover, these results align with the observation that the two tagged colonies, where most colony members were tagged, did not initiate tandem runs (Table 1).

**Fig 3.**
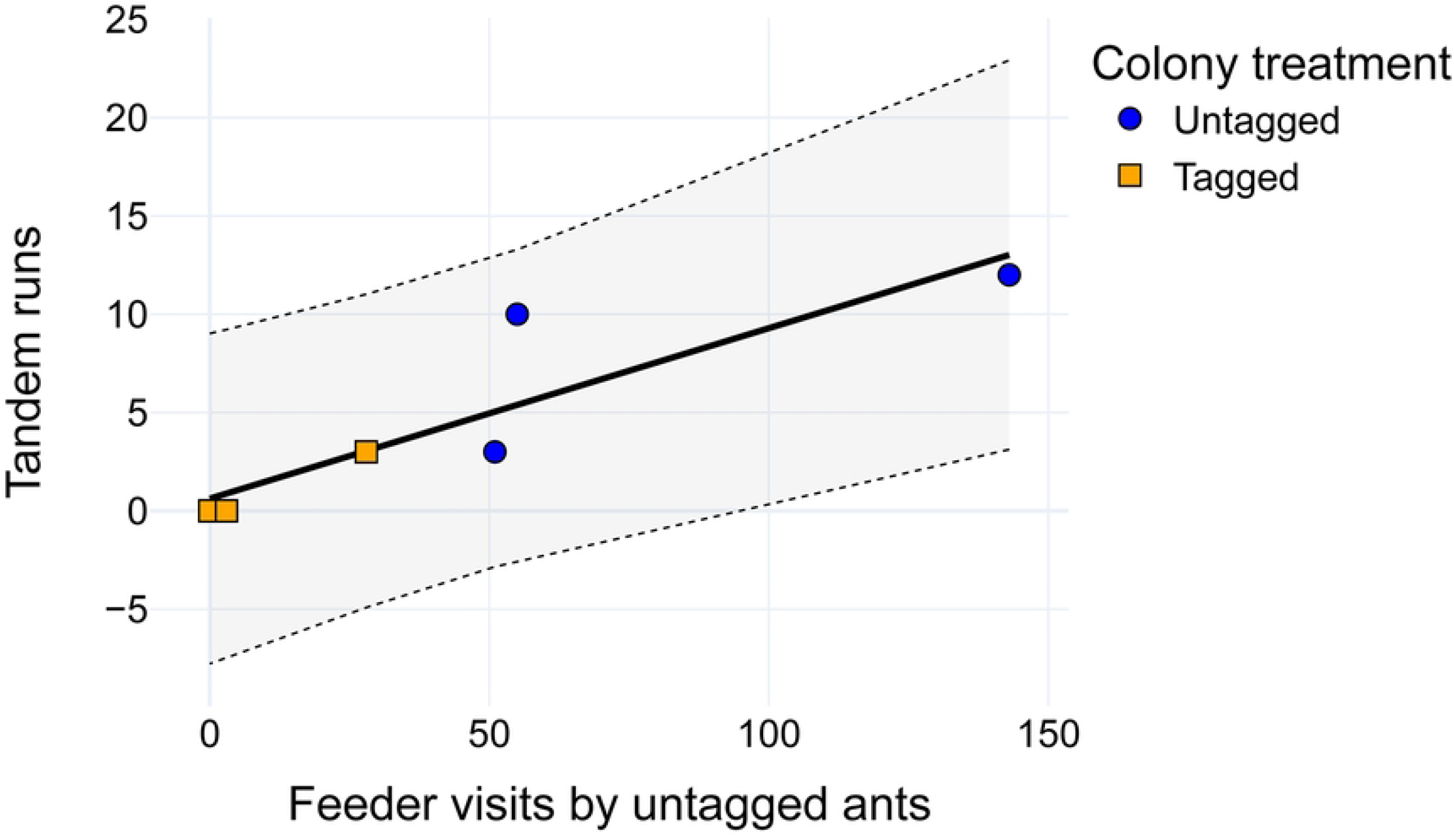
Significant positive relationship between feeder visits by untagged ants and tandem runs in tagged and untagged colonies: OLS regression analysis reveals a significant positive relationship between the number of feeder visits by untagged ants and the number of tandem runs led within tagged (orange, square) and untagged (blue, circle) colonies (y = 0.09x + 0.62, R^2^=0.79, Adjusted-R^2^=0.74, *P* = 0.02). The solid line represents the regression line, while the light grey bands between the dashed lines indicate the 95% confidence interval.

## Discussion

Our study demonstrates that tracking tags, an emerging method for exploring collective behavior in social insects [16-18, 35-39], can have negative effects on ant behavior. Within partially tagged colonies, we find that tags did not reduce individual foraging activity, with tagged ants visiting the food source as frequently as untagged ants. This finding aligns with previous observations that tracking tags present no immediate debilitating effects on movement [17]. However, our analysis revealed a marked difference in recruitment behavior, with tagged ants participating in tandem running significantly less than untagged ants. Using untagged colonies as a control group, we can rule out potential influences from arena design factors such as light levels and food source distance. Furthermore, the comparison between tagged and untagged colonies supports the detrimental effects of tags on foraging, with tagged colonies participating in fewer tandem runs overall.

### Why did tagged ants not recruit nestmates?

Notably, we only observed tagged ants participating as tandem run followers, never as leaders. The absence of tagged ants initiating recruitment is not likely due to the added weight of the tags since ants can readily transport weights exceeding their own [40, 41]. Additionally, in our experiments, we observed tagged ants carrying dead ants to graveyards, indicating that the weight of the tags did not impede their carrying abilities. Alternatively, if tagged ants were insensitive or inaccessible to antennal contact, it could prevent them from leading tandem runs but not participating as followers. Since tandem running relies on the tactile contact between the follower’s antennae and the leader’s gaster, if their gasters were damaged during the tagging process or from the materials used (*e*.*g*., adhesives), it could reduce the leaders’ ability to detect the followers’ antennae. Similarly, reduced agility from the tagging process could limit their coordination ability. Since tandem-running followers regulate the movement of tandem pairs [42], a speed mismatch might explain our results for tagged and untagged pairs; however, it would not explain the absence of tagged pairs tandem-running. Additional experiments are necessary to pinpoint the exact cause of reduced recruitment by tagged ants.

One potential solution to minimize the impact of tracking tags on ant behavior is to reduce the size of the tags. Previous *Temnothorax* studies that used smaller devices (0.5 mm^2^) found no evidence of device attachment impacting behavior (AprilTags [18] and RFID transponders [43, 44]). However, reducing the tag size further (<0.8 mm^2^) presents practical challenges and increased costs. Tagged-based tracking systems require high-quality input images to detect and identify tags, which entails significant experimental control and investment. For instance, manufacturing miniature tags requires specialized printing to achieve the necessary edge definition (square shape) for computer readability [8], which can be expensive. Furthermore, the quality of optics and lighting limits the success of these systems [16]. As a result, automated tracking setups typically use high-resolution (≥ 4K) cameras [18, 38, 45-47] and high-end optics [16] to capture tags at the requisite tracking size (pixels/mm). Lastly, to extend the trackable area captured by a single sensor, researchers have implemented multiple camera/sensor setups [16, 45]. However, these setups necessitate additional investments in equipment such as cameras, storage, and processing power (CPUs & GPUs). In short, an additional reduction in tag size would require a costly investment in modifying the tracking system.

Our study highlights the importance of implementing controls beyond visual assessments to detect the behavioral impacts of tracking devices. While several studies have examined the impact of tracking devices on animal well-being [48], many devices, particularly the newer ones, have yet to be thoroughly investigated. With the prevalence and ongoing development of miniaturized tracking technology [49], it is crucial to establish limits and species-specific protocols for tracking devices. Future studies using tracking devices should carefully consider the natural history of their study organism [37] and incorporate control groups into their experimental designs [50]. Although tracking devices are powerful tools for understanding complex behavioral patterns, it is crucial to carefully consider their potential impact on animal behavior to ensure accurate conclusions.

## Acknowledgments

We thank Richard Hall and Andy Davis for their insightful comments on the manuscript. This work was supported by the National Science Foundation (#2118012) and James L. Carmon Scholarship.

## Supporting information

S1 Table 1. Correlations between colony variables and tandem running events.

## References

1. Couzin ID, Heins C. Emerging technologies for behavioral research in changing environments. Trends in Ecology & Evolution. 2022.

2. Westley PA, Berdahl AM, Torney CJ, Biro D. Collective movement in ecology: from emerging technologies to conservation and management. The Royal Society; 2018. p. 20170004.

3. Panadeiro V, Rodriguez A, Henry J, Wlodkowic D, Andersson M. A review of 28 free animal-tracking software applications: current features and limitations. Lab Animal. 2021;50(9):246–54. Epub 20210729. doi: 10.1038/s41684-021-00811-1. PubMed PMID: 34326537.

4. Dell AI, Bender JA, Branson K, Couzin ID, De Polavieja GG, Noldus LPJJ, et al. Automated image-based tracking and its application in ecology. Trends in Ecology & Evolution. 2014;29(7):417–28. Epub 20140605. doi: 10.1016/j.tree.2014.05.004. PubMed PMID: 24908439.

5. Cagnacci F, Boitani L, Powell RA, Boyce MS. Animal ecology meets GPS-based radiotelemetry: a perfect storm of opportunities and challenges. Philosophical Transactions of the Royal Society B: Biological Sciences. 2010;365(1550):2157–62.

6. Kays R, Crofoot MC, Jetz W, Wikelski M. Terrestrial animal tracking as an eye on life and planet. Science. 2015;348(6240):aaa2478. doi: doi:10.1126/science.aaa2478.

7. Thorup K, Holland RA. The bird GPS – long-range navigation in migrants. Journal of Experimental Biology. 2009;212(22):3597–604. doi: 10.1242/jeb.021238.

8. Graving JM, Chae D, Naik H, Li L, Koger B, Costelloe BR, et al. DeepPoseKit, a software toolkit for fast and robust animal pose estimation using deep learning. eLife. 2019;8. Epub 20191001. doi: 10.7554/elife.47994. PubMed PMID: 31570119; PubMed Central PMCID: PMCPMC6897514.

9. Wilson RP. The price tag. Nature. 2011;469(7329):164–5. doi: 10.1038/469164a.

10. Gillies N, Fayet AL, Padget O, Syposz M, Wynn J, Bond S, et al. Short-term behavioural impact contrasts with long-term fitness consequences of biologging in a long-lived seabird. Scientific Reports. 2020;10(1):15056. doi: 10.1038/s41598-020-72199-w.

11. Aldridge HDJN, Brigham RM. Load Carrying and Maneuverability in an Insectivorous Bat: a Test of the 5% “Rule” of Radio-Telemetry. Journal of Mammalogy. 1988;69(2):379–82. doi: 10.2307/1381393.

12. Mellor DJ, Beausoleil NJ, Stafford KJ. Marking amphibians, reptiles and marine mammals: animal welfare, practicalities and public perceptions in New Zealand: Department of Conservation Wellington, New Zealand; 2004.

13. Fiala M, editor Comparing ARTag and ARToolkit Plus fiducial marker systems. IEEE International Workshop on Haptic Audio Visual Environments and their Applications; 2005: IEEE.

14. Garrido-Jurado S, Muñoz-Salinas R, Madrid-Cuevas FJ, Medina-Carnicer R. Generation of fiducial marker dictionaries using Mixed Integer Linear Programming. Pattern Recognition. 2016;51:481–91. doi: 10.1016/j.patcog.2015.09.023.

15. Olson E, editor AprilTag: A robust and flexible visual fiducial system. 2011 IEEE international conference on robotics and automation; 2011: IEEE.

16. Mersch DP, Crespi A, Keller L. Tracking Individuals Shows Spatial Fidelity Is a Key Regulator of Ant Social Organization. Science. 2013;340(6136):1090–3. Epub 20130418. doi: 10.1126/science.1234316. PubMed PMID: 23599264.

17. Stroeymeyt N, Grasse AV, Crespi A, Mersch DP, Cremer S, Keller L. Social network plasticity decreases disease transmission in a eusocial insect. Science. 2018;362(6417):941–5. doi: 10.1126/science.aat4793. PubMed PMID: 30467168.

18. Richardson TO, Stroeymeyt N, Crespi A, Keller L. Two simple movement mechanisms for spatial division of labour in social insects. Nature Communications. 2022;13(1):6985. doi: 10.1038/s41467-022-34706-7.

19. Franks NR, Pratt SC, Mallon EB, Britton NF, Sumpter DJ. Information flow, opinion polling and collective intelligence in house-hunting social insects. Philos Trans R Soc Lond B Biol Sci. 2002;357(1427):1567–83. doi: 10.1098/rstb.2002.1066. PubMed PMID: 12495514; PubMed Central PMCID: PMCPMC1693068.

20. Sasaki T, Pratt SC. The Psychology of Superorganisms: Collective Decision Making by Insect Societies. Annual Review of Entomology. 2018;63(1):259–75. Epub 20171004. doi: 10.1146/annurev-ento-020117-043249. PubMed PMID: 28977775.

21. Pratt S, Mallon E, Sumpter D, Franks N. Quorum sensing, recruitment, and collective decision-making during colony emigration by the ant Leptothorax albipennis. Behavioral Ecology and Sociobiology. 2002;52(2):117–27. doi: 10.1007/s00265-002-0487-x.

22. Shaffer Z, Sasaki T, Pratt SC. Linear recruitment leads to allocation and flexibility in collective foraging by ants. Animal Behaviour. 2013;86(5):967–75.

23. Mallon EB, Pratt SC, Franks NR. Individual and collective decision-making during nest site selection by the ant Leptothorax albipennis. Behavioral Ecology and Sociobiology. 2001;50(4):352–9. doi: 10.1007/s002650100377.

24. Möglich M. Social organization of nest emigration in Leptothorax (Hym., Form.). Insectes Sociaux. 1978;25(3):205–25. doi: 10.1007/BF02224742.

25. Franklin EL, Richardson TO, Sendova-Franks AB, Robinson EJ, Franks NR. Blinkered teaching: tandem running by visually impaired ants. Behavioral Ecology and Sociobiology. 2011;65(4):569–79.

26. Bhatkar A, Whitcomb W. Artificial diet for rearing various species of ants. Florida Entomologist. 1970:229–32.

27. Graving J. pinpoint: behavioral tracking using 2D barcode tags v0. 0.1-alpha. Version v0 01-alpha-DOI doi. 2017;105281.

28. Pratt SC, Brooks SE, Franks NR. The Use of Edges in Visual Navigation by the Ant Leptothorax albipennis. Ethology. 2001;107(12):1125–36. doi: 10.1046/j.1439-0310.2001.00749.x.

29. Sasaki T, Danczak L, Thompson B, Morshed T, Pratt SC. Route learning during tandem running in the rock ant Temnothorax albipennis. Journal of Experimental Biology. 2020;223(9):jeb221408. Epub 20200515. doi: 10.1242/jeb.221408. PubMed PMID: 32414865.

30. Tomar S. Converting video formats with FFmpeg. Linux Journal. 2006;2006(146):10.

31. Hörl D, Rojas Rusak F, Preusser F, Tillberg P, Randel N, Chhetri RK, et al. BigStitcher: reconstructing high-resolution image datasets of cleared and expanded samples. Nature methods. 2019;16(9):870–4.

32. Bradski G. The OpenCV Library. Dr Dobb’s Journal: Software Tools for the Professional Programmer. 2000;25(11):120–3.

33. Virtanen P, Gommers R, Oliphant TE, Haberland M, Reddy T, Cournapeau D, et al. SciPy 1.0: fundamental algorithms for scientific computing in Python. Nature methods. 2020;17(3):261–72.

34. Seabold S, Perktold J, editors. Statsmodels: Econometric and statistical modeling with python. Proceedings of the 9th Python in Science Conference; 2010: Austin, TX.

35. Richardson TO, Kay T, Braunschweig R, Journeau OA, Rüegg M, McGregor S, et al. Ant behavioral maturation is mediated by a stochastic transition between two fundamental states. Current Biology. 2021;31(10):2253–60.e3. doi: https://doi.org/10.1016/j.cub.2020.05.038.

36. Boenisch F, Rosemann B, Wild B, Dormagen D, Wario F, Landgraf T. Tracking all members of a honey bee colony over their lifetime using learned models of correspondence. Frontiers in Robotics and AI. 2018;5:35.

37. Crall JD, Gravish N, Mountcastle AM, Combes SA. BEEtag: A Low-Cost, Image-Based Tracking System for the Study of Animal Behavior and Locomotion. PLOS ONE. 2015;10(9):e0136487. doi: 10.1371/journal.pone.0136487.

38. Quque M, Bles O, Bénard A, Héraud A, Meunier B, Criscuolo F, et al. Hierarchical networks of food exchange in the black garden ant Lasius niger. Insect science. 2021;28(3):825–38.

39. Crall JD, Souffrant AD, Akandwanaho D, Hescock SD, Callan SE, Coronado WM, et al. Social context modulates idiosyncrasy of behaviour in the gregarious cockroach Blaberus discoidalis. Animal Behaviour. 2016;111:297–305.

40. Hölldobler B, Wilson EO. The ants: Belknap, Harvard Univ Press,; 1990.

41. Traniello J. Comparative foraging ecology of north temperate ants: the role of worker size and cooperative foraging in prey selection. Insectes Sociaux. 1987;34(2):118–30.

42. Valentini G, Masuda N, Shaffer Z, Hanson JR, Sasaki T, Walker SI, et al. Division of labour promotes the spread of information in colony emigrations by the ant Temnothorax rugatulus. Proceedings of the Royal Society B: Biological Sciences. 2020;287(1924):20192950. Epub 20200401. doi: 10.1098/rspb.2019.2950. PubMed PMID: 32228408; PubMed Central PMCID: PMCPMC7209055.

43. Robinson EJH, Smith FD, Sullivan KME, Franks NR. Do ants make direct comparisons? Proceedings of the Royal Society B: Biological Sciences. 2009;276(1667):2635–41. doi: 10.1098/rspb.2009.0350.

44. Robinson EJH, Richardson TO, Sendova-Franks AB, Feinerman O, Franks NR. Radio tagging reveals the roles of corpulence, experience and social information in ant decision making. Behavioral Ecology and Sociobiology. 2009;63(5):627–36. doi: 10.1007/s00265-008-0696-z.

45. Greenwald EE, Baltiansky L, Feinerman O. Individual crop loads provide local control for collective food intake in ant colonies. eLife. 2018;7:e31730. doi: 10.7554/eLife.31730.

46. Greenwald E, Segre E, Feinerman O. Ant trophallactic networks: simultaneous measurement of interaction patterns and food dissemination. Scientific Reports. 2015;5(1):12496. doi: 10.1038/srep12496.

47. Richardson TO, Liechti JI, Stroeymeyt N, Bonhoeffer S, Keller L. Short-term activity cycles impede information transmission in ant colonies. PLOS Computational Biology. 2017;13(5):e1005527. Epub 20170510. doi: 10.1371/journal.pcbi.1005527. PubMed PMID: 28489896; PubMed Central PMCID: PMCPMC5443549.

48. Murray DL, Fuller MR. A Critical Review of the Effects of Marking on the Biology of Vertebrates. In: Pearl MC, Boitani L, Fuller TK, editors. Research Techniques in Animal Ecology. Controversies and Consequences. 2 ed: Columbia University Press; 2000. p. 15–64.

49. Krause J, Krause S, Arlinghaus R, Psorakis I, Roberts S, Rutz C. Reality mining of animal social systems. Trends in Ecology & Evolution. 2013;28(9):541–51. doi: https://doi.org/10.1016/j.tree.2013.06.002.

50. Balmori A. Radiotelemetry and wildlife: Highlighting a gap in the knowledge on radiofrequency radiation effects. Science of The Total Environment. 2016;543:662–9. doi: https://doi.org/10.1016/j.scitotenv.2015.11.073.

